# OptiClust: Improved method for assigning amplicon-based sequence data to operational taxonomic units

**DOI:** 10.1101/096537

**Authors:** Sarah L. Westcott, Patrick D. Schloss

## Abstract

Assignment of 16S rRNA gene sequences to operational taxonomic units (OTUs) is a computational bottleneck in the process of analyzing microbial communities. Although this has been an active area of research, it has been difficult to overcome the time and memory demands while improving the quality of the OTU assignments. Here we developed a new OTU assignment algorithm that iteratively reassigns sequences to new OTUs to optimize the Matthews correlation coefficient (MCC), a measure of the quality of OTU assignments. To assess the new algorithm, OptiClust, we compared it to ten other algorithms using 16S rRNA gene sequences from two simulated and four natural communities. Using the OptiClust algorithm, the MCC values averaged 15.2 and 16.5% higher than the OTUs generated when we used the average neighbor and distance-based greedy clustering with VSEARCH, respectively. Furthermore, on average, OptiClust was 94.6-times faster than the average neighbor algorithm and just as fast as distance-based greedy clustering with VSEARCH. An empirical analysis of the efficiency of the algorithms showed that the time and memory required to perform the algorithm scaled quadratically with the number of unique sequences in the dataset. The significant improvement in the quality of the OTU assignments over previously existing methods will significantly enhance downstream analysis by limiting the splitting of similar sequences into separate OTUs and merging of dissimilar sequences into the same OTU. The development of the OptiClust algorithm represents a significant advance that is likely to have numerous other applications.

## Importance

The analysis of microbial communities from diverse environments using 16S rRNA gene sequencing has expanded our knowledge of the biogeography of microorganisms. An important step in this analysis is the assignment of sequences into taxonomic groups based on their similarity to sequences in a database or based on their similarity to each other, irrespective of a database. In this study, we present a new algorithm for the latter approach. The algorithm, OptiClust, seeks to optimize a metric of assignment quality by shuffling sequences between taxonomic groups. We found that OptiClust produces more robust assignments and does so in a rapid and memory efficient manner. This advance will allow for a more robust analysis of microbial communities and the factors that shape them.

## Introduction

Amplicon-based sequencing has provided incredible insights into Earth’s microbial biodiversity (1, 2). It has become common for studies to include sequencing millions of 16S rRNA gene sequences across hundreds of samples (3, 4). This is three to four orders of magnitude greater sequencing depth than was previously achieved using Sanger sequencing (5, 6). The increased sequencing depth has revealed novel taxonomic diversity that is not adequately represented in reference databases (1, 3). However, the advance has forced re-engineering of methods to overcome the rate and memory limiting steps in computational pipelines that process raw sequences through the generation of tables containing the number of sequences in different taxa for each sample (7–10). A critical component to these pipelines has been the assignment of amplicon sequences to taxonomic units that are ether defined based on similarity to a reference or operationally based on the similarity of the sequences to each other within the dataset (11, 12).

A growing number of algorithms have been developed to cluster sequences into OTUs. These algorithms can be classified into three general categories. The first category of algorithms has been termed closed-reference or phylotyping (13, 14). Sequences are compared to a reference collection and clustered based on the reference sequences that they are similar to. This approach is fast; however, the method struggles when a sequence is similar to multiple reference sequences that may have different taxonomies and when it is not similar to sequences in the reference (15). The second category of algorithms has been called *de novo* because they assign sequences to OTUs without the use of a reference (14). These include hierarchical algorithms such as nearest, furthest, and average neighbor (16) and algorithms that employ heuristics such as abundance or distance-based greedy clustering as implemented in USEARCH (17) or VSEARCH (18), Sumaclust, OTUCLUST (19), and Swarm (20). *De novo* methods are agglomerative and tend to be more computationally intense. It has proven difficult to know which method generates the best assignments. A third category of algorithm is open reference clustering, which is a hybrid approach (3, 14). Here sequences are assigned to OTUs using closed-reference clustering and sequences that are not within a threshold of a reference sequence are then clustered using a *de novo* approach. This category blends the strengths and weaknesses of the other method and adds the complication that closed-reference and *de novo* clustering use different OTU definitions. These three categories of algorithms take different approaches to handling large datasets to minimize the time and memory requirements while attempting to assign sequences to meaningful OTUs.

Several metrics have emerged for assessing the quality of OTU assignment algorithms. These have included the time and memory required to run the algorithm (3, 20–22), agreement between OTU assignments and the sequences’ taxonomy (20, 22–32), sensitivity of an algorithm to stochastic processes (33), the number of OTUs generated by the algorithm (23, 34), and the ability to regenerate the assignments made by other algorithms (3, 35). Unfortunately, these methods fail to directly quantify the quality of the OTU assignments. An algorithm may complete with minimal time and memory requirements or generate an idealized number of OTUs, but the composition of the OTUs could be incorrect. These metrics also tend to be subjective. For instance, a method may appear to recapitulate the taxonomy of a synthetic community with known taxonomic structure, but do a poor job when applied to real communities with poorly defined taxonomic structure or for sequences that are prone to misclassification. As an alternative, we developed an approach to objectively benchmark the clustering quality of OTU assignments (13, 15, 36). This approach counts the number of true positives (TP), true negatives (TN), false positives (FP), and false negatives (FN) based on the pairwise distances. Sequence pairs that are within the user-specified threshold and are clustered together represent TPs and those in different OTUs are FNs. Those sequence pairs that have a distance larger than the threshold and are not clustered in the same OTU are TNs and those in the same OTU are FPs. These values can be synthesized into a single correlation coefficient, the Matthews correlation coefficient (MCC), which measures the correlation between observed and predicted classifications and is robust to cases where there is an uneven distribution across the confusion matrix (37). Consistently, the average neighbor algorithm was identified as among the best or the best algorithm. Other hierarchical algorithms such as furthest and nearest neighbor, which do not permit the formation of FPs or FNs, respectively, fared significantly worse. The distance-based greedy clustering as implemented in VSEARCH has also performed well. The computational resources required to complete the average neighbor algorithm can be significant for large datasets and so there is a need for an algorithm that efficiently produces consistently high quality OTU assignments.

These benchmarking efforts have assessed the quality of the clusters after the completion of the algorithm. In the current study we developed and benchmarked a new *de novo* clustering algorithm that uses real time calculation of the MCC to direct the progress of the clustering. The result is the OptiClust algorithm, which produces significantly better sequence assignments while making efficient use of computational resources.

## Results

### OptiClust algorithm

The OptiClust algorithm uses the pairs of sequences that are within a desired threshold of each other (e.g. 0.03), a list of all sequence names in the dataset, and the metric that should be used to assess clustering quality. A detailed description of the algorithm is provided for a toy dataset in the Supplementary Material. Briefly, the algorithm starts by placing each sequence either within its own OTU or into a single OTU. The algorithm proceeds by interrogating each sequence and re-calculating the metric for the cases where the sequence stays in its current OTU, is moved to each of the other OTUs, or is moved into a new OTU. The location that results in the best clustering quality indicates whether the sequence should remain in its current OTU or be moved to a different or new OTU. Each iteration consists of interrogating every sequence in the dataset. Although numerous options are available for optimizing the clusters and for assessing the quality of the clusters within the mothur-based implementation of the algorithm (e.g. sensitivity, specificity, accuracy, F1-score, etc.), the default metric for optimization and assessment is MCC because it includes all four parameters from the confusion matrix (Figure S1; Table S1). The algorithm continues until the optimization metric stabilizes or until it reaches a defined stopping criteria.

### OptiClust-generated OTUs are more robust than those from other methods

To evaluate the OptiClust algorithm and compare its performance to other algorithms, we utilized six datasets including two synthetic communities and four previously published large datasets generated from soil, marine, human, and murine samples (Table 1). When we seeded the OptiClust algorithm with each sequence in a separate OTU and ran the algorithm until complete convergence, the MCC values averaged 15.2 and 16.5% higher than the OTUs using average neighbor and distance-based greedy clustering (DGC) with VSEARCH, respectively (Figure 1; Table S1). The number of OTUs formed by the various methods was negatively correlated with their MCC value (*ρ* =-0.47; p<0.001). The OptiClust algorithm was considerably faster than the hierarchical algorithms and somewhat slower than the heuristic-based algorithms. Across the six datasets, the OptiClust algorithm was 94.6-times faster than average neighbor and just as fast as DGC with VSEARCH. The human dataset was a challenge for a number of the algorithms. OTUCLUST and SumaClust were unable to cluster the human dataset in less than 50 hours and the average neighbor algorithm required more than 45 GB of RAM. The USEARCH-based methods were unable to cluster the human data using the 32-bit free version of the software that limits the amount of RAM to approximately 3.5 GB. These data demonstrate that OptiClust generated significantly more robust OTU assignments than existing methods across a diverse collection of datasets with performance that was comparable to popular methods.

**Table 1.** Description of datasets used to evaluate the OptiClust algorithm and compare its performance to other algorithms. Each dataset contains sequences from the V4 region of the16S rRNA gene. The number of distances for each dataset are those that were less than or equal to 0.03. The number of OTUs were determined using the OptiClust algorithm. The even and staggered datasets were generated by extracting the V4 region from full length reference sequences and the datasets from the natural communities were generated by sequencing the V4 region using a Illumina MiSeq with either paired 150 or 250 nt reads.

| Dataset (Ref.) | Read Length | Samples | Total Seqs. | Unique Seqs. | Distances | OTUs |
| --- | --- | --- | --- | --- | --- | --- |
| Soil (41) | 150 | 18 | 948,243 | 143,677 | 11,775,167 | 40,216 |
| Marine (42) | 250 | 7 | 1,384,988 | 75,923 | 12,908,857 | 25,787 |
| Mice (40) | 250 | 360 | 2,825,495 | 32,447 | 6,988,306 | 2,658 |
| Human (39) | 250 | 489 | 20,951,841 | 121,281 | 38,544,315 | 11,648 |
| Even (34, 36) | NA | NA | 1,155,800 | 11,558 | 29,694 | 7,651 |
| Staggered (34, 36) | NA | NA | 1,156,550 | 11,558 | 29,694 | 7,653 |

**Figure 1.**
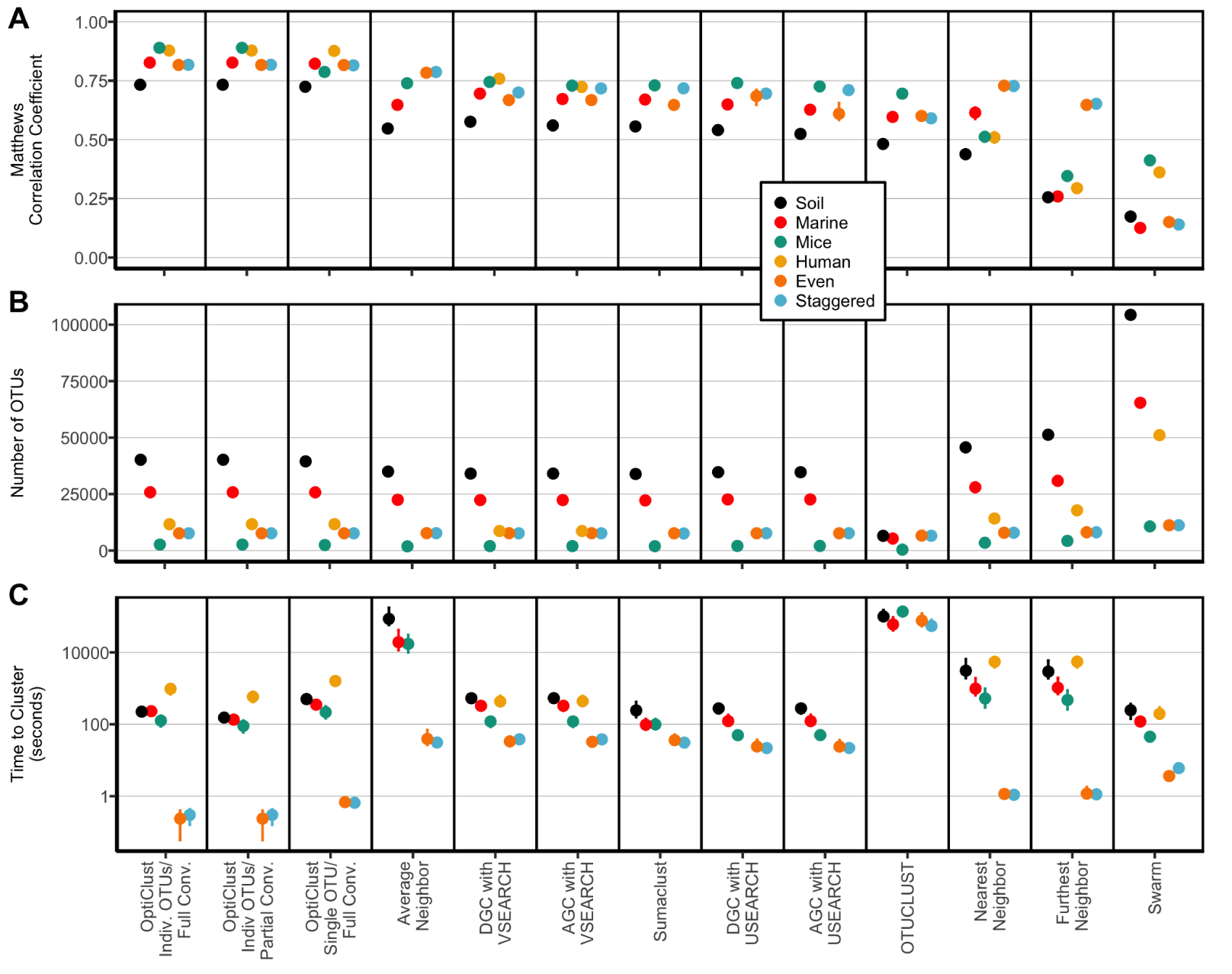
Comparison of de novo clustering algorithms. Plot of MCC (A), number of OTUs (B), and execution times (C) for the comparison of *de novo* clustering algorithms when applied to four natural and two synthetic datasets. The first three columns of each figure contain the results of clustering the datasets (i) seeding the algorithm with one sequence per OTU and allowing the algorithm to proceed until the MCC value no longer changed; (ii) seeding the algorithm with one sequence per OTU and allowing the algorithm to proceed until the MCC changed by less than 0.0001; (iii) seeding the algorithm with all of the sequences in one OTU and allowing the algorithm to proceed until the MCC value no longer changed. The human dataset could not be clustered by the average neighbor, Sumaclust, USEARCH, or OTUCLUST with less than 45 GB of RAM or 50 hours of execution time. The median of 10 re-orderings of the data is presented for each method and dataset. The range of observed values is indicated by the error bars, which are typically smaller than the plotting symbol.

### OptiClust stopping criteria

By default, the mothur-based implementation of the algorithm stops when the optimization metric changes by less than 0.0001; however, this can be altered by the user. This implementation also allows the user to stop the algorithm if a maximum number of iterations is exceeded. By default mothur uses a maximum value of 100 iterations. The justification for allowing incomplete convergence was based on the observation that numerous iterations are performed that extend the time required to complete the clustering with minimal improvement in clustering (Figure S2). We evaluated the results of clustering to partial convergence (i.e. a change in the MCC value that was less than 0.0001) or until complete convergence of the MCC value (i.e. until it did not change between iterations) when seeding the algorithm with each sequence in a separate OTU (Figure 1). The small difference in MCC values between the output from partial and complete convergence resulted in a difference in the median number of OTUs that ranged between 1.5 and 17.0 OTUs. This represented a difference of less than 0.15%. Among the four natural datasets, between 3 and 6 were needed to achieve partial convergence and between 8 and 12 iterations were needed to reach full convergence. The additional steps required between 1.4 and 1.7 times longer to complete the algorithm. These results suggest that achieving full convergence of the optimization metric adds computational effort; however, considering full convergence took between 2 and 17 minutes the extra effort was relatively small. Although the mothur’s default setting is partial convergence, the remainder of our analysis used complete convergence to be more conservative.

### Effect of seeding OTUs on OptiClust performance

By default the mothur implementation of the OptiClust algorithm starts with each sequence in a separate OTU. An alternative approach is to start with all of the sequences in a single OTU. We found that the MCC values for clusters generated seeding OptiClust with the sequences as a single OTU were between 0 and 11.5% lower than when seeding the algorithm with sequences in separate OTUs (Figure 1). Interestingly, with the exception of the human dataset (0.2% more OTUs), the number of OTUs was as much as 7.0% lower (mice) than when the algorithm was seeded with sequence in separate OTUs. Finally, the amount of time required to cluster the data when the algorithm was seeded with a single OTU was between 1.5 and 2.9-times longer than if sequences were seeded as separate OTUs. This analysis demonstrates that seeding the algorithm with sequences as separate OTUs resulted in the best OTU assignments in the shortest amount of time.

### OptiClust-generated OTUs are as stable as those from other algorithms

One concern that many have with *de novo* clustering algorithms is that their output is sensitive to the initial order of the sequences because each algorithm must break ties where a sequence could be assigned to multiple OTUs. An additional concern specific to the OptiClust algorithm is that it may stabilize at a local optimum. To evaluate these concerns we compared the results obtained using ten randomizations of the order that sequences were given to the algorithm. The median coefficient of variation across the six datasets for MCC values obtained from the replicate clusterings using OptiClust was 0.1% (Figure 1). We also measured the coefficient of variation for the number of OTUs across the six datasets for each method. The median coefficient of variation for the number of OTUs generated using OptiClust was 0.1%. Confirming our previous results (15), all of the methods we tested were stable to stochastic processes. Of the methods that involved randomization, the coefficient of variation for MCC values was considerably smaller with OptiClust than the other methods and the coefficient of variation for the number of OTUs was comparable to the other methods. The variation observed in clustering quality suggested that the algorithm does not appear to converge to a locally optimum MCC value. More importantly, the random variation does yield output of a similarly high quality.

### Time and memory required to complete Optimization-based clustering scales efficiently

Although not as important as the quality of clustering, the amount of time and memory required to assign sequences to OTUs is a legitimate concern. We observed that the time required to complete the OptiClust algorithm (Figure 1C) paralleled the number of pairwise distances that were smaller than 0.03 (Table 1). To further evaluate how the speed and memory usage scaled with the number of sequences in the dataset, we measured the time required and maximum RAM usage to cluster 20, 40, 60, 80, and 100% of the unique sequences from each of the natural datasets using the OptiClust algorithm (Figure 2). Within each iteration of the algorithm, each sequence is compared to every other sequence and each comparison requires a recalculation of the confusion matrix. This would result in a worst case algorithmic complexity on the order of N^3^, where N is the number of unique sequences. Because the algorithm only needs to keep track of the sequence pairs that are within the threshold of each other, it is likely that the implementation of the algorithm is more efficient. To empirically determine the algorithmic complexity, we fit a power law function to the data in Figure 2A. We observed power coefficients between 1.7 and 2.5 for the marine and human datasets, respectively. The algorithm requires storing a matrix that contains the pairs of sequences that are close to each other as well as a matrix that indicates which sequences are clustered together. The memory required to store these matrices is on the order of N^2^, where N is the number of unique sequences. In fact, when we fit a power law function to the data in Figure 2B, the power coefficients were 1.9. Using the four natural community datasets, doubling the number of sequences in a dataset would increase the time required to cluster the data by 4 to 8-fold and increase the RAM required by 4-fold. It is possible that future improvements to the implementation of the algorithm could improve this performance.

**Figure 2.**
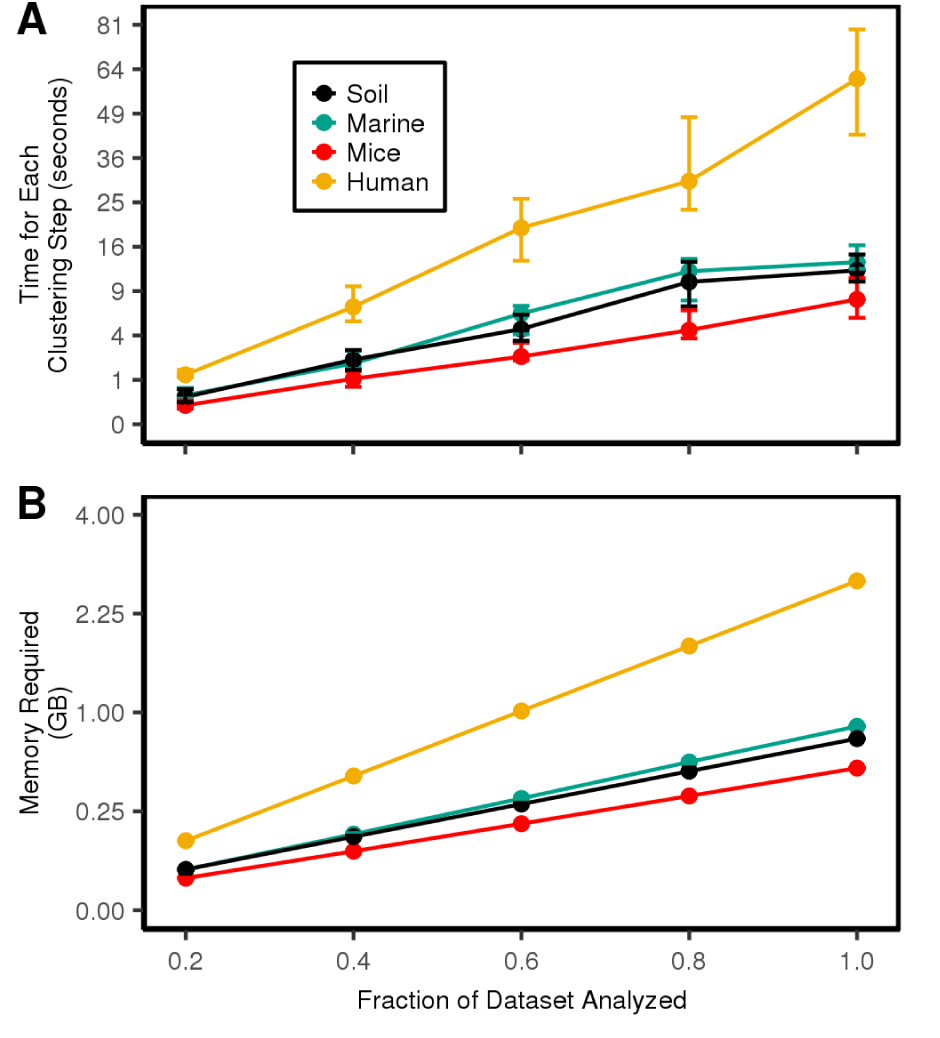
OptiClust performance. The average execution time (A) and memory usage (B) requiredto cluster the four natural datasets. The confidence intervals indicate the range between the minimum and maximum values. The y-axis is scaled by the square root to demonstrate the relationship between the time and memory requirements relative to the number of unique sequences squared.

### Cluster splitting heuristic generates OTUs that are as good as non-split approach

We previously described a heuristic to accelerate OTU assignments where sequences were first classified to taxonomic groups and within each taxon sequences were assigned to OTUs using the average neighbor clustering algorithm (13). This method is similar to open reference clustering except that in our approach all sequences are subjected to *de novo* clustering following classification whereas in open reference clustering only those sequences that cannot be classified are subjected to *de novo* clustering. Our cluster splitting approach accelerated the clustering and reduced the memory requirements because the number of unique sequences was effectively reduced by splitting sequences across taxonomic groups. Furthermore, because sequences in different taxonomic groups are assumed to belong to different OTUs they are independent, which permits parallelization and additional reduction in computation time. Reduction in clustering quality is encountered in this approach if there are errors in classification or if two sequences within the desired threshold belong to different taxonomic groups. It is expected that these errors would increase as the taxonomic level goes from kingdom to genus. To characterize the clustering quality, we classified each sequence at each taxonomic level and calculated the MCC values using OptiClust, average neighbor, and DGC with VSEARCH when splitting at each taxonomic level (Figure 3). For each method, the MCC values decreased as the taxonomic resolution increased; however, the decrease in MCC was not as large as the difference between clustering methods. As the resolution of the taxonomic levels increased, the clustering quality remained high, relative to clusters formed from the entire dataset (i.e. kingdom-level). The MCC values when splitting the datasets at the class and genus levels were within 98.0 and 93.0%, respectively, of the MCC values obtained from the entire dataset. These decreases in MCC value resulted in the formation of as many as 4.7 and 22.5% more OTUs, respectively, than were observed from the entire dataset. These errors were due to the generation of additional false negatives due to splitting similar sequences into different taxonomic groups. For the datasets included in the current analysis, the use of the cluster splitting heuristic was probably not worth the loss in clustering quality. However, as datasets become larger, it may be necessary to use the heuristic to clustering the data into OTUs.

**Figure 3.**
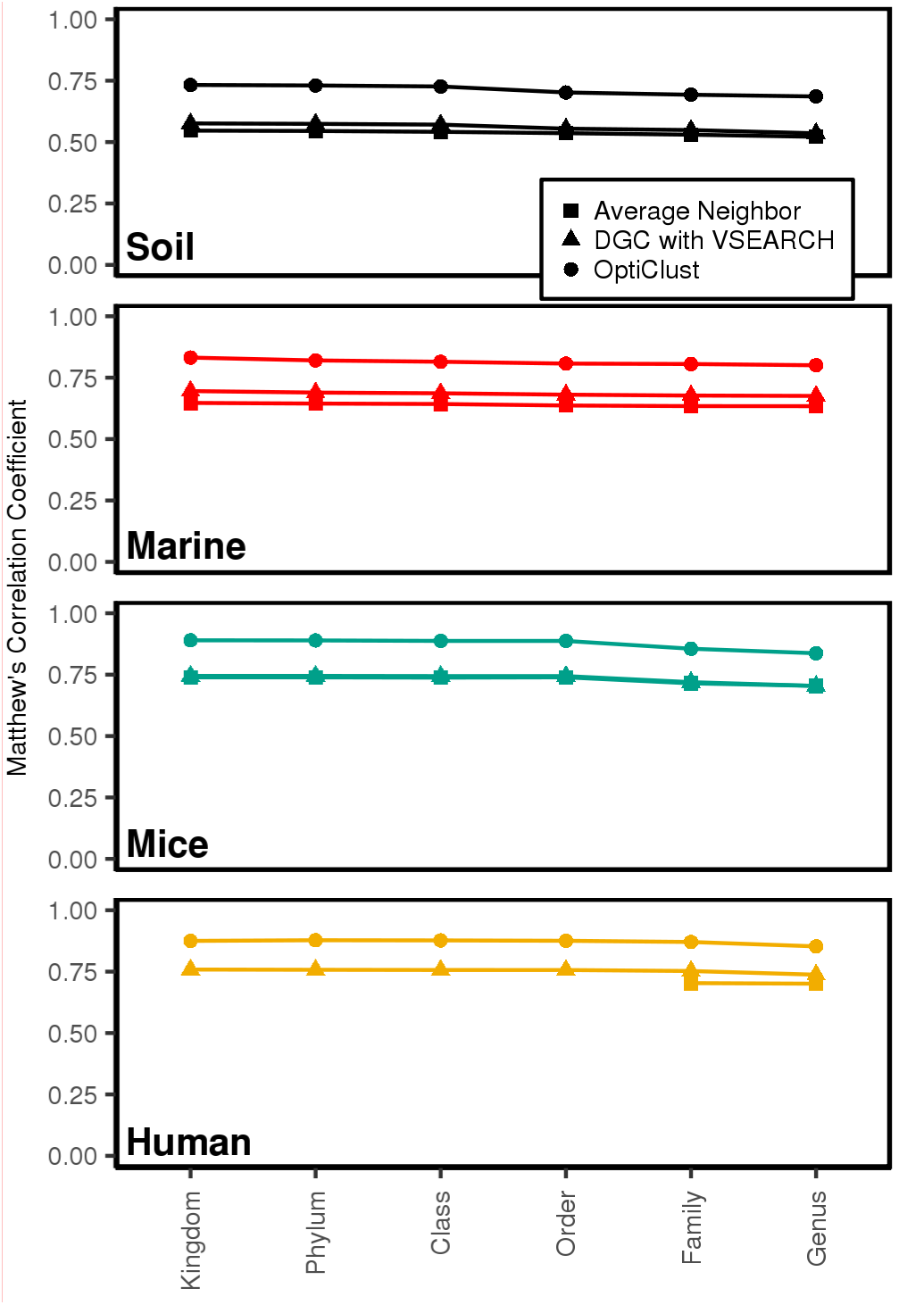
Effects of taxonomically splitting the datasets on clustering quality. The datasets were split at each taxonomic level based on their classification using a naive Bayesian classifier and clustered using average neighbor, VSEARCH-based DGC, and OptiClust.

## Discussion

Myriad methods have been proposed for assigning 16S rRNA gene sequences to OTUs. Each claim improved performance based on speed, memory usage, representation of taxonomic information, and number of OTUs. Each of these metrics is subjective and do not actually indicate the quality of the clustering. This led us to propose using the MCC as a metric for assessing the quality of clustering, post hoc. Here, we described a new clustering method that seeks to optimize clustering based on an objective criterion that measures clustering quality in real time. In the OptiClust algorithm, clustering is driven by optimizing a metric that assesses whether any two sequences should be grouped into the same OTU. The result is clusters that are significantly more robust and is efficient in the time and memory required to cluster the sequences into OTUs. This makes it more tractable to analyze large datasets without sacrificing clustering quality as was previously necessary using heuristic methods.

The cluster optimization procedure is dependent on the metric that is chosen for optimization. We employed the MCC because it includes the four values from a confusion matrix. Other algorithms such as the furthest neighbor and nearest neighbor algorithms minimize the number of FP and FN, respectively; however, these suffer because the number of FN and FP are not controlled, respectively (13, 16). Alternatively, one could optimize based on the sensitivity, specificity, or accuracy, which are each based on two values from the confusion matrix or they could optimize based on the F1-score, which is based on three values from the confusion matrix. Because these metrics do not balance all four parameters equally, it is likely that one parameter will dominate in the optimization procedure. For example, optimizing for sensitivity could lead to a large number of FPs. More FPs increases the number of OTUs while more FNs collapses OTUs together. It is difficult to know which is worse since community richness and diversity are linked to the number of OTUs. In addition, increasing the number of FNs would overstate the differences between communities while increasing the number of FPs would overstate their similarity. Therefore, it is important to jointly minimize the number of FPs and FNs. With this in mind, we decided to optimize utilizing the MCC. It is possible that other metrics that balance the four parameters could be developed and employed for optimization of the clustering.

The OptiClust algorithm is relatively simple. For each sequence it effectively asks whether the MCC value will increase if the sequence is moved to a different OTU including creating a new OTU. If the value does not change, it remains in the current OTU. The algorithm repeats until the MCC value stabilizes. Assuming that the algorithm is seeded with each sequence in a separate OTU, it does not appear that the algorithm converges to a local optimum. Furthermore, execution of the algorithm with different random number generator seeds produces OTU assignments of consistently high quality. Future improvements to the implementation of the algorithm could provide optimization to further improve its speed and susceptibility to find a local optimum. Users are encouraged to repeat the OTU assignment several times to confirm that they have found the best OTU assignments.

Our previous MCC-based analysis of clustering algorithms indicated that the average neighbor algorithm consistently produced the best OTU assignments with the DGC-based method using USEARCH also producing robust OTU assignments. The challenge in using the average neighbor algorithm is that it requires a large amount of RAM and is computationally demanding. This led to the development of a splitting approach that divides the clustering across distinct taxonomic groups (13). The improved performance provided by the OptiClust algorithm likely makes such splitting unnecessary for most current datasets. We have demonstrated that although the OTU assignments made at the genus level are still better than that of other methods, the quality is not as good as that found without splitting. The loss of quality is likely due to misclassification because of limitations in the clustering algorithms and reference databases. The practical significance of such small differences in clustering quality remain to be determined; however, based on the current analysis, it does appear that the number of OTUs is artificially inflated. Regardless, the best clustering quality should be pursued given the available computer resources.

The time and memory required to execute the OptiClust algorithm scaled proportionally to the number of unique sequences raised to the second power. The power for the time requirement is affected by the similarity of the sequences in the dataset with datasets containing more similar sequences having a higher power. Also, the number of unique sequences is the basis for both the amount of time and memory required to complete the algorithm. Both the similarity of sequences and number of unique sequences can be driven by the sequencing error since any errors will increase the number of unique sequences and these sequences will be closely related to the perfect sequence. This underscores the importance of reducing the noise in the sequence data (7). If sequencing errors are not remediated and are relatively randomly distributed, then it is likely that the algorithm will require an unnecessary amount of time and RAM to complete.

The rapid expansion in sequencing capacity has demanded that the algorithms used to assign 16S rRNA gene sequences to OTUs be efficient while maintaining robust assignments. Although database-based approaches have been proposed to facilitate this analysis, they are limited by their limited coverage of bacterial taxonomy and by the inconsistent process used to name taxa. The ability to assign sequences to OTUs using an algorithm that optimizes clustering by directly measuring quality will significantly enhance downstream analysis. The development of the OptiClust algorithm represents a significant advance that is likely to have numerous other applications.

## Materials and Methods

### Sequence data and processing steps

To evaluate the OptiClust and the other algorithms we created two synthetic sequence collections and four sequence collections generated from previously published studies. The V4 region of the 16S rRNA gene was used from all datasets because it is a popular region that can be fully sequenced with two-fold coverage using the commonly used MiSeq sequencer from Illumina (7). The method for generating the simulated datasets followed the approach used by Kopylova et al. (34) and Schloss (36). Briefly, we randomly selected 10,000 uniques V4 fragments from 16S rRNA gene sequences that were unique from the SILVA non-redundant database (38). A community with an even relative abundance profile was generated by specifying that each sequence had a frequency of 100 reads. A community with a staggered relative abundance profile was generated by specifying that the abundance of each sequence was a randomly drawn integer sampled from a uniform distribution between 1 and 200. Sequence collections collected from human feces (39), murine feces (40), soil (41), and seawater (42) were used to characterize the algorithms’ performance with natural communities. These sequence collections were all generated using paired 150 or 250 nt reads of the V4 region. We re-processed all of the reads using a common analysis pipeline that included quality score-based error correction (7), alignment against a SILVA reference database (38, 43), screening for chimeras using UCHIME (9), and classification using a naive Bayesian classifier with the RDP training set requiring an 80% confidence score (10).

### Implementation of clustering algorithms

In addition to the OptiClust algorithm we evaluated ten different *de novo* clustering algorithms. These included three hierarchical algorithms, average neighbor, nearest neighbor, and furthest neighbor, which are implemented in mothur (v.1.39.0) (11). Seven heuristic methods were also used including abundance-based greedy clustering (AGC) and (distance-based greedy clustering) DGC as implemented in USEARCH (v.6.1) (17) and VSEARCH (v.2.3.3) ((18)], OTUCLUST (v.0.1) (19), SumaClust (v.1.0.20), and Swarm (v.2.1.9) (20). With the exception of Swarm each of these methods uses distance-based thresholds to report OTU assignments. We also evalauted the ability of OptiClust to optimize to metrics other than MCC. These included accuracy, F1-score, negative predictive value, positive predictive value, false discovery rate, senitivity, specificity, the sum of TPs and TNs, the sum of FPs and FNs, and the number of FNs, FPs, TNs, and TPs (Figure S1; Table S1).

### Benchmarking

We evaluated the quality of the sequence clustering, reproducibility of the clustering, the speed of clustering, and the amount of memory required to complete the clustering. To assess the quality of the clusters generated by each method, we counted the cells within a confusion matrix that indicated how well the clusterings represented the distances between the pair of sequences (13). Pairs of sequences that were in the same OTU and had a distance less than 3% were true positives (TPs), those that were in different OTUs and had a distance greater than 3% were true negatives (TNs), those that were in the same OTU and had a distance greater than 3% were false positives (FPs), and those that were in different OTUs and had a distance less than 3% were false negatives (FNs). To synthesize the matrix into a single metric we used the Matthews correlation coefficient using the sens.spec command in mothur using the following equations.

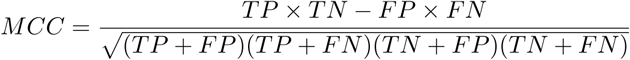

To assess the reproducibility of the algorithms we randomized the starting order of each sequence collection ten times and ran each algorithm on each randomized collection. We then measured the MCC for each randomization and quantified their percent coefficient of variation (% CV; 100 times the ratio of the standard deviation to the mean).

To assess how the the memory and time requirements scaled with the number of sequences included in each sequence collection, we randomly subsampled 20, 40, 60, or 80% of the unique sequences in each collection. We obtained 10 subsamples at each depth for each dataset and ran each collection (N= 50 = 5 sequencing depths × 10 replicates) through each of the algorithms. We used the timeout script to quantify the maximum RAM used and the amount of time required to process each sequence collection (https://github.com/pshved/timeout). We limited each algorithm to 45 GB of RAM and 50 hours using a single processor.

### Data and code availability

The workflow utilized commands in GNU make (v.3.81), GNU bash (v.4.1.2), mothur (v.1.39.0) (11), and R (v.3.3.2) (44). Within R we utilized the wesanderson (v.0.3.2) (45), dplyr (v.0.5.0) (46), tidyr (v.0.6.0) (47), cowplot (v.0.6.3) (48), and ggplot2 (v.2.2.0.9000) (49) packages. A reproducible version of this manuscript and analysis is available at https://github.com/SchlossLab/Westcott_OptiClust_mSphere_2017.

## Acknowledgements

This work was supported through funding from the National Institutes of Health to PDS (P30DK034933). SLW designed, implemented, and evaluated the algorithm. PDS designed and evaluated the algorithm. Both authors wrote and edited the manuscript.

---

**Supplemental text**. Worked example of how OptiClust algorithm clusters sequences into OTUs.

**Figure S1.**
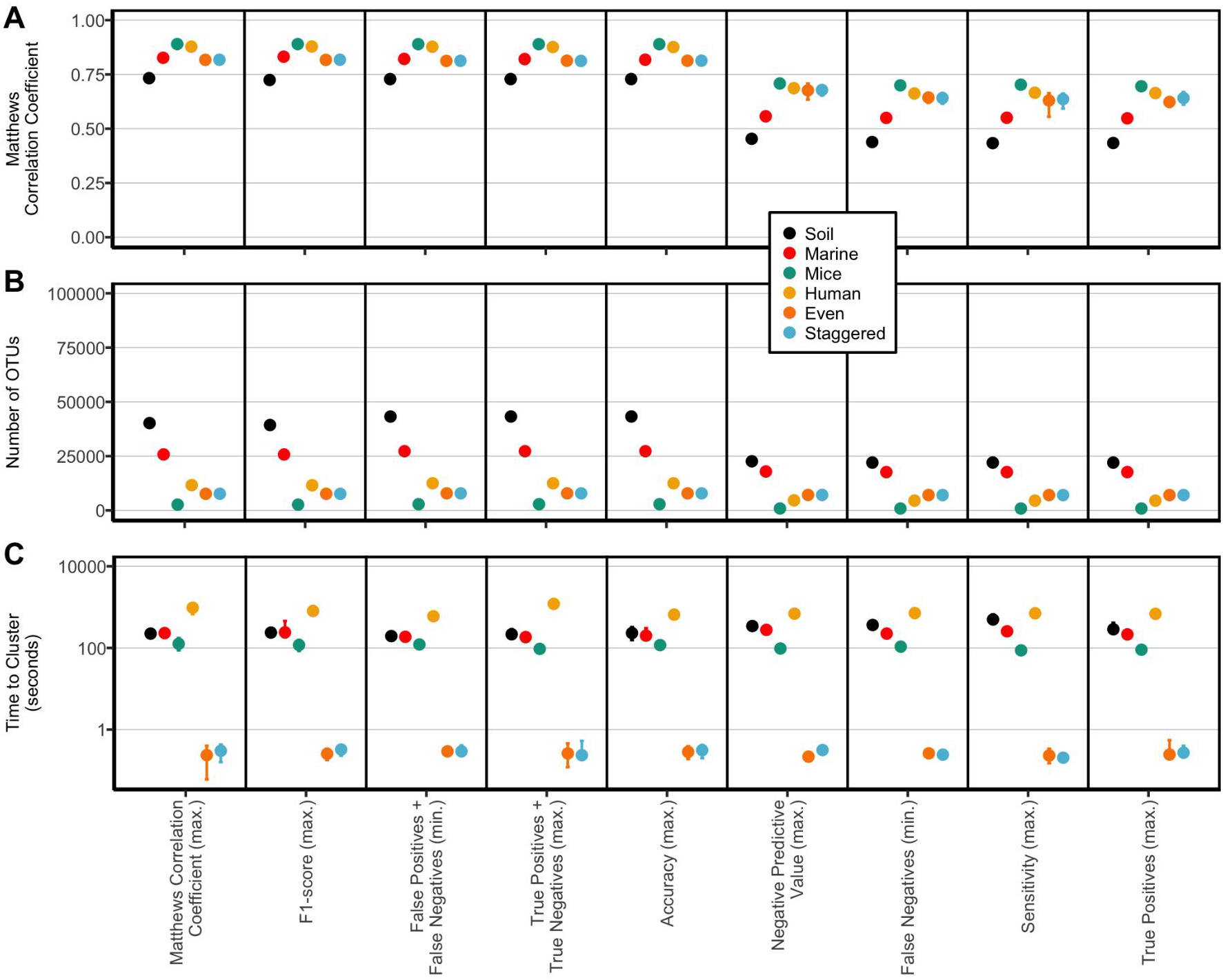
The OptiClust algorithm is able to effectively cluster sequences into OTUs by minimizing or maximizing numerous metrics. Plot of MCC (A), number of OTUs (B), and execution times (C) for the comparison of output from the OptiClust algorithm when to minimizing or maximizing a variety of parameters when applied to four natural and two synthetic datasets. Within mothur, OTU assignments can also be made using other metrics including minimizing false positives and maximizing the specificity, positive predictive value, and true negatives; however, these all resulted in sequences being assigned to separate OTUs, which resulted in no false positives and the maximum number of true negatives. The error bars indicate the range of values observed for 10 replicates.

**Figure S2.**
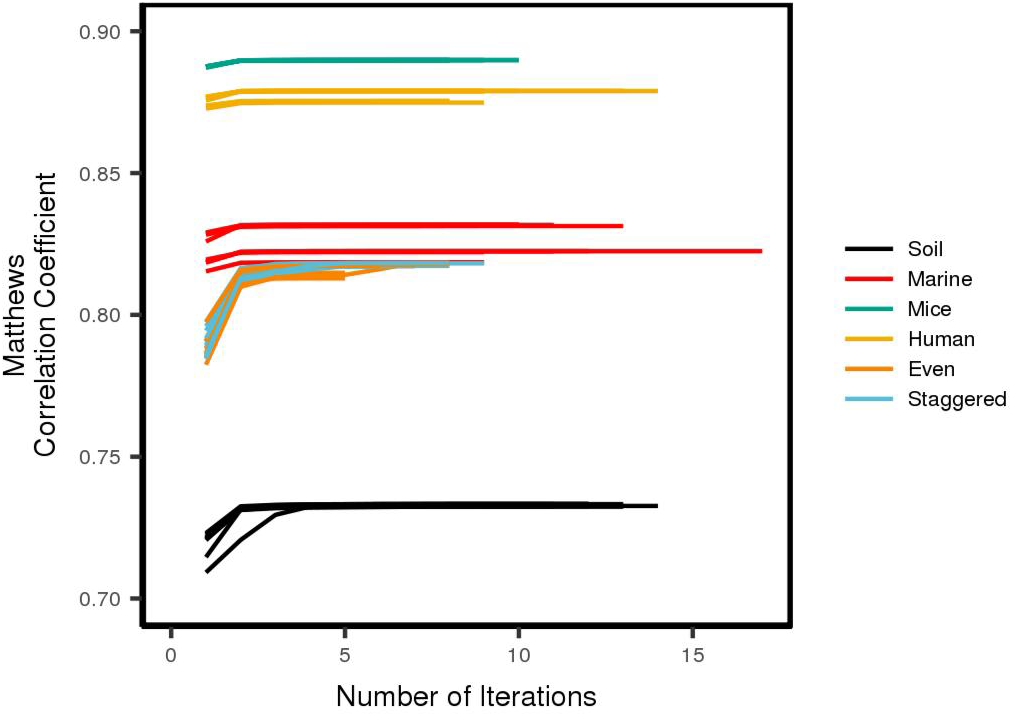
The OptiClust algorithm rapidly converges to optimize the Matthews correlation coefficient. The six datasets were clustered into OTUs using the OptiClust algorithm seeking to maximize the Matthews correlation coefficient. This was repeated 10 times for each dataset.

**Table S1.**
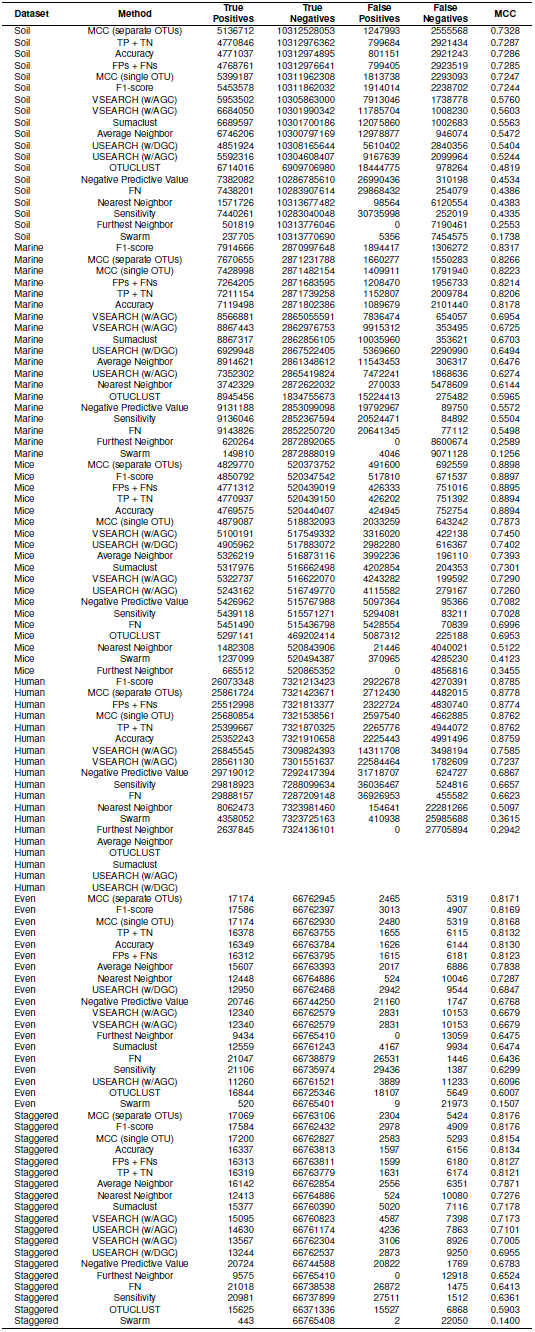
Summary of the average number of true positives, true negatives, false positives, false negatives and the resulting Matthews correlation coefficient for each of the clustering methods that were analyzed in this study for each of the six datasets. Blank values indicate that those conditions could not be completed in 50 hours with 45 GB of RAM.

## Supplemental Text

To provide a detailed demonstration of how the OptiClust algorithm works consider a relatively simple example where there are 50 sequences. After aligning the sequences and calculating the pairwise distances between the sequences there are 15 pairs of sequences with a distance below the desired threshold of 0.03:

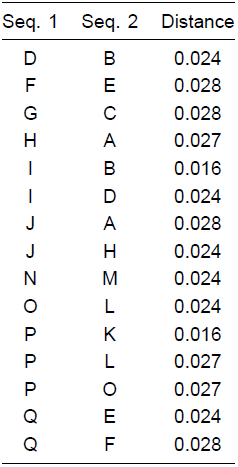

The other 1210 distances were larger than 0.03 and are not needed for the analysis because they are taken to be true negatives. For example, the distance between sequences A and B is larger than 0.03. So, if they are in different OTUs then that would be a true negative (TN) and if they are in the same OTU then that would be a false positive (FP). Alternatively, because sequences B and D are closer to each other than 0.03 (i.e. 0.024) if they are in separate OTUs then that would be a false negative (FN) and if they are in the same OTU then that would be a true positive (TP). It is important to note that the algorithm assumes that there are no duplicate sequences and the actual abundance is saved elsewhere to be substituted later when counting the frequency distribution of each OTU across the samples included in the analysis.

The algorithm starts by seeding sequences either into individual OTUs or into a single OTU. As demonstrated in Figure 1, seeding the sequences into randomly ordered individual OTUs generates better results and is faster than starting with a single OTU. Among the 15 pairwise distances that are smaller than 0.03, there are 17 sequences that are labeled A through Q. A separate pool is created for the 33 other sequences that are not within 0.03 of any other sequence and are thus to be placed into 33 separate OTUs. In the diagrams below, this pool is designated as “…”. Having seeded the initial OTUs there are 0 TPs, 1210 TNs, 0 FPs, and 15 FNs. Initially the number of FNs corresponds to the number of distances less than 0.03, the number of TNs is the number of total distances (i.e. 1225) minus the number of distances less than 0.03. The number of TPs, TNs, FPs, and FNs should sum to the total number of distances. The resulting Matthew’s Correlation Coefficient (MCC) is 0.00. The algorithm next goes through each sequence sequentially to determine whether the MCC value would be increased by removing it from its current OTU to join other sequences in a new OTU or to create its own OTU.

The demonstration of the algorithm starts with sequence A. Notice that it is within 0.03 of sequences H and J. There are three options: sequence A could remain as its own OTU, it could join with sequence H, or it could join with sequence J.

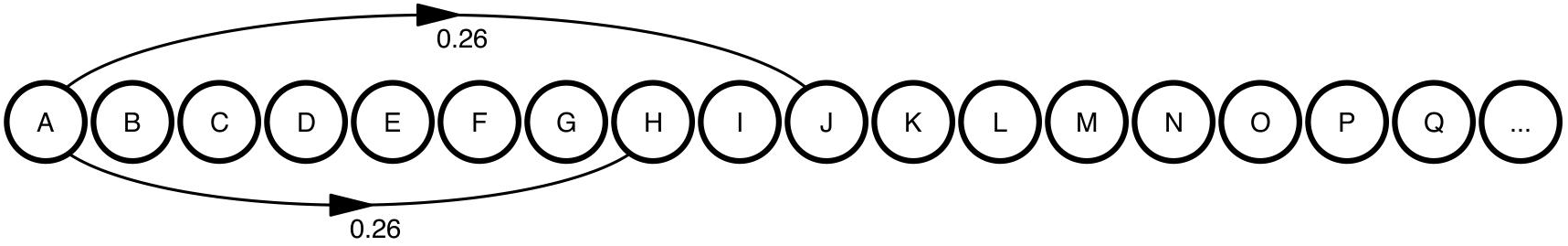

If it stays as its own OTU, the MCC value would remain 0.00. If it joined with sequences H or J the number of TPs would increase by 1 and the number of FNs would decrease by 1. In either case the MCC would be 0.26. In this case, joining H or J would result in an improved MCC and so the algorithm randomly selects which sequences to join. For this demonstration, it will form a new OTU with sequence H. This results in 1 TPs, 1210 TNs, 0 FPs, 14 FNs, and an MCC of 0.26.

Sequence B is processed by the same process as sequence A. Sequence B is within 0.03 of sequences D and I. Again, there are three options: sequence B could remain as its own OTU, it could join with sequence D, or it could join with sequence I.

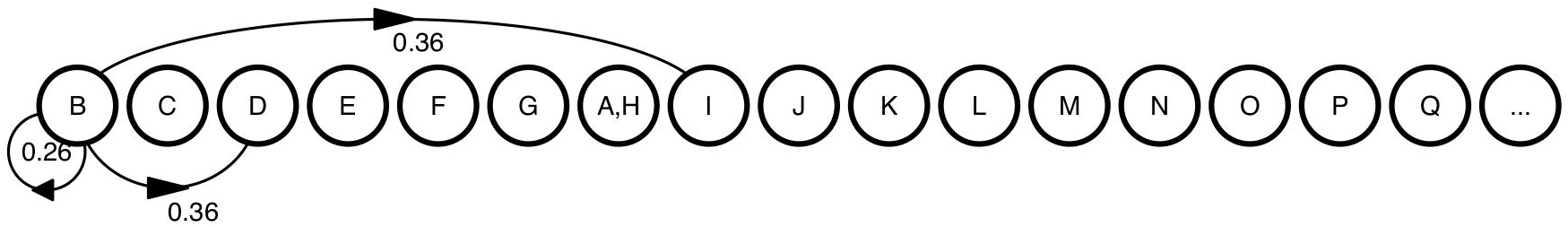

If it remains as its own OTU, the MCC value would remain 0.26. If it joined with sequences D or I the number of TPs would increase by 1 and the number of FNs would decrease by 1. In either case the MCC would be 0.36. Joining D or I would result in an improved MCC and so the algorithm randomly selects which sequences to join. For this demonstration it will form a new OTU with sequence D. This results in 2 TPs, 1210 TNs, 0 FPs, 13 FNs, and an MCC of 0.36.

Sequence C is within 0.03 of sequence G creating two options: remain as its own OTU or join with sequence G.

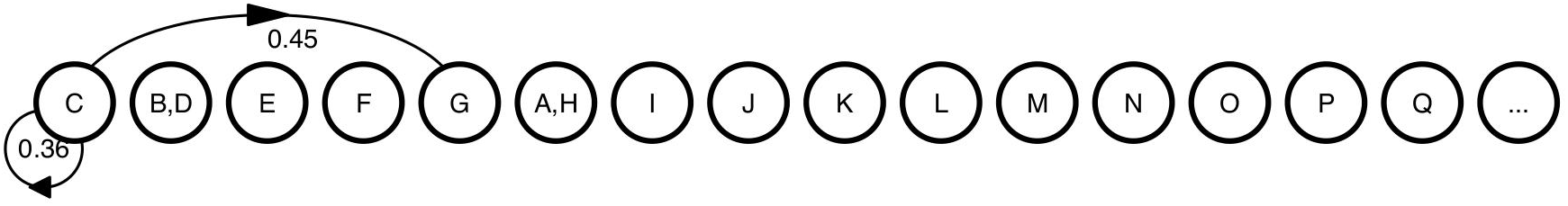

If it remains in its own OTU, the MCC value would remain 0.36. If it joined with sequence G the number of TPs would increase by 1 and the number of FNs would decrease by 1 resulting in an MCC value of 0.45. Because of the improved MCC value, sequence C joins with sequence G to form a new OTU. This results in 3 TPs, 1210 TNs, 0 FPs, 12 FNs, and an MCC of 0.45.

For sequence D the algorithm gets more complicated since sequence D is already in an OTU with sequence B. As seen when considering sequence B, sequence D is within 0.03 of sequence B and it is also similar to sequence I. Now there are three options: sequence D could remain with sequence B in an OTU, it could leave that OTU and join with sequence I, or it could form a new OTU where it is the sole member.

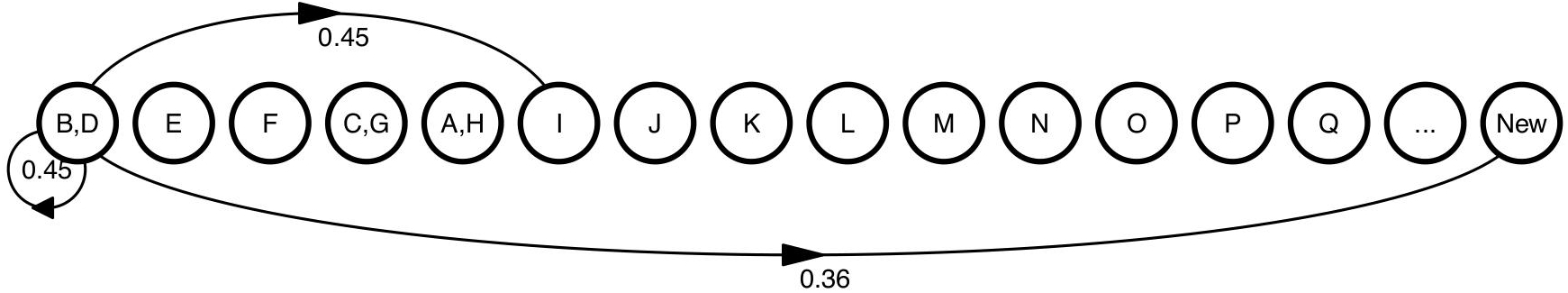

If sequence D remains with B, the MCC value would remain 0.45. If it joined with sequence I the number of TPs and FNs would stay constant resulting in an MCC value of 0.51. If it formed a new OTU by itself, the number of TPs would decrease by one and the number of FNs would increase by 1 resulting in an MCC value of 0.36. Because the MCC values for staying in the OTU with B or leaving the OTU to join the OTU with I are the same, the algorithm would again randomly chose between the two options. For demonstration purposes, sequence D will remain in its OTU with sequence B. This results in no changes in the four parameters or the MCC value.

For sequence E, the same type of options are available as when sequences A and B were processed. Sequence E could remain on its own or it could join with sequences F or Q.

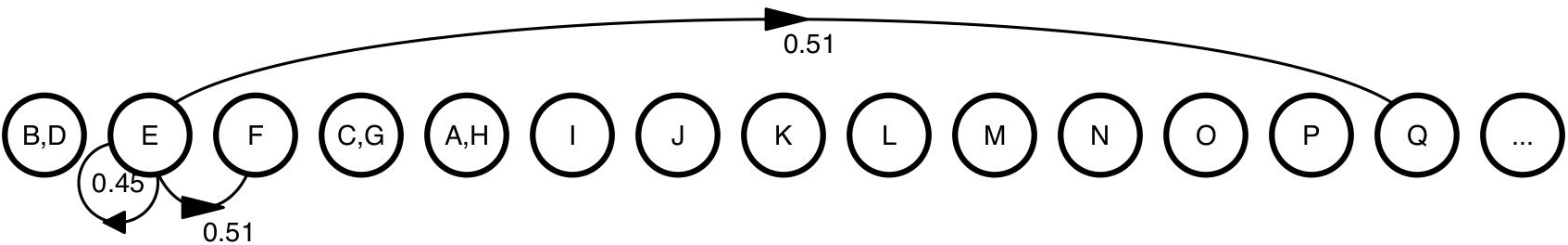

Similar to the earlier cases, the MCC value for joining another sequence is larger than staying on its own. Because the MCC values for joining F or Q are the same, the algorithm randomly selects which sequences to join. For this demonstration sequence E will form a new OTU with sequence F. This results in 4 TPs, 1210 TNs, 0 FPs, 11 FNs, and an MCC of 0.51.

For sequence F, the steps taken by the algorithm are the same as earlier for sequence D. Here, sequence F could remain in an OTU with sequence E, it could leave and form an OTU with sequence Q, or it could form a new OTU on its own.

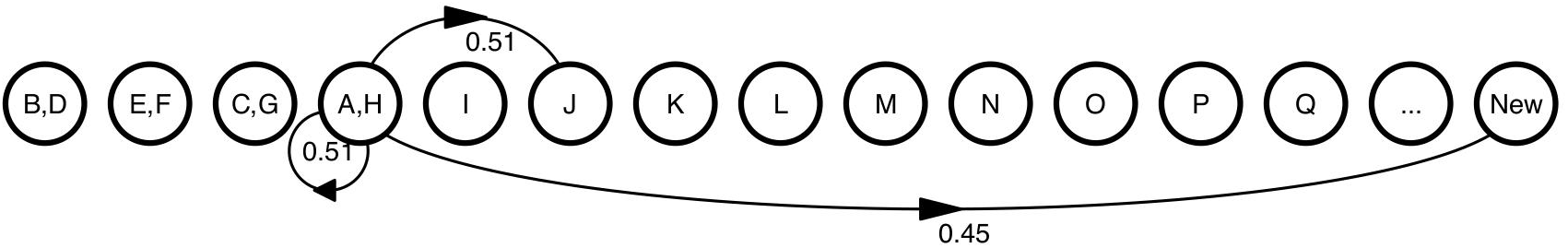

Again, the MCC value for sequence F remaining with sequence E is the same as for leaving to form an OTU with sequence Q. Both options are superior to leaving to form a new OTU on its own. The algorithm randomly chooses between the two options. For demonstration purposes, sequence F will remain in its OTU with sequence E. This results in no changes in the four parameters or the MCC value.

The decisions for sequence G are similar to sequence F, with the exception that the only choices are to stay in an OTU with another sequence (C) or to form a new OTU on its own.

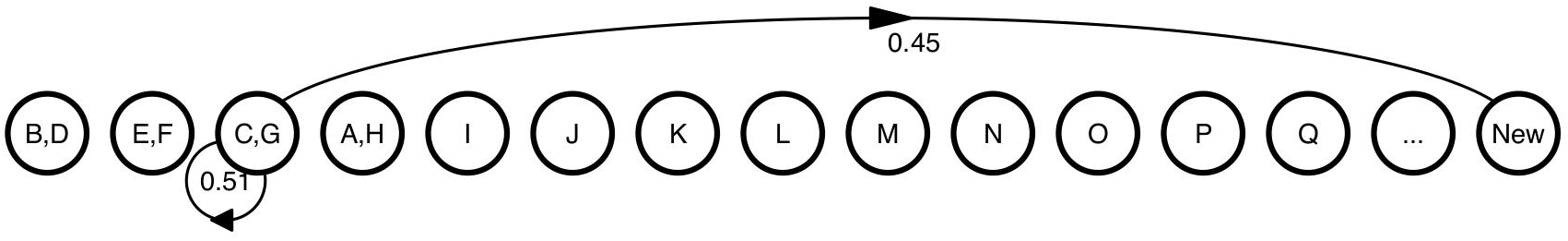

Leaving to form a new OTU results in a lower MCC value and so the algorithm leaves sequence G with sequence C. This results in no changes in the four parameters or the MCC value.

For sequence H the steps taken are the same as seen earlier for sequence B.

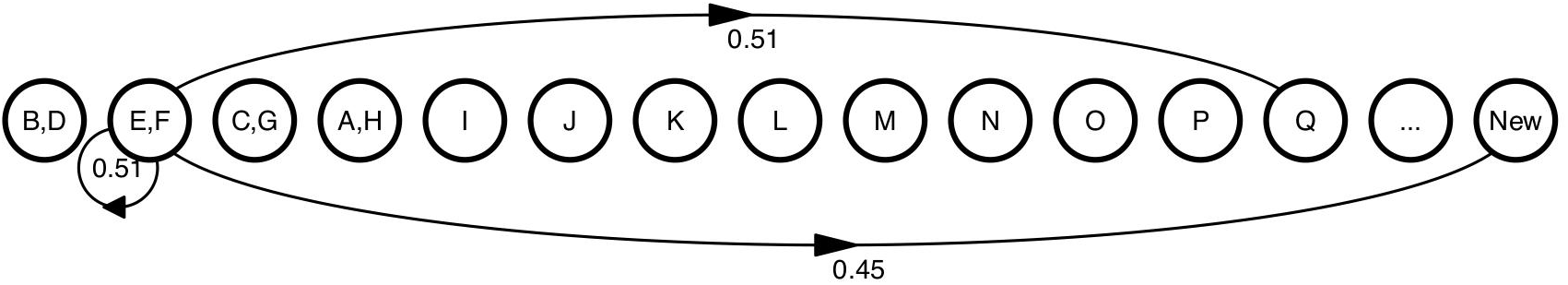

For demonstration purposes sequence H will remain in its OTU with sequence A.

Moving to sequence I, the process is similar to what was done earlier with sequences A and B. The only difference is that because sequence I is similar to both sequences B and D the increase in TP and decrease in FN will be double.

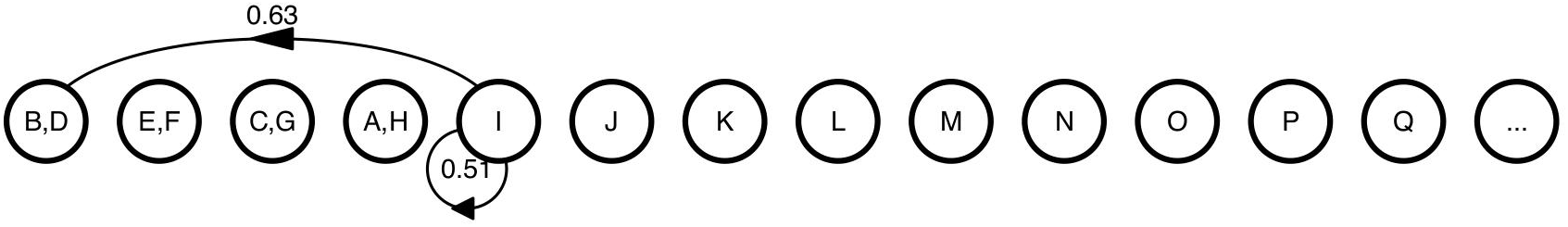

If sequence I remains in an OTU by itself, the MCC value will be 0.51. If it joined the OTU with sequences B and D, then the number of TPs would increase by 2 and the number of FNs would decrease by 2 resulting in an improved MCC value of 0.63. This is the choice that is taken resulting in 6 TPs, 1210 TNs, 0 FPs, and 9 FNs.

Processing sequence J is the same as sequence I since sequence J is close to both A and H.

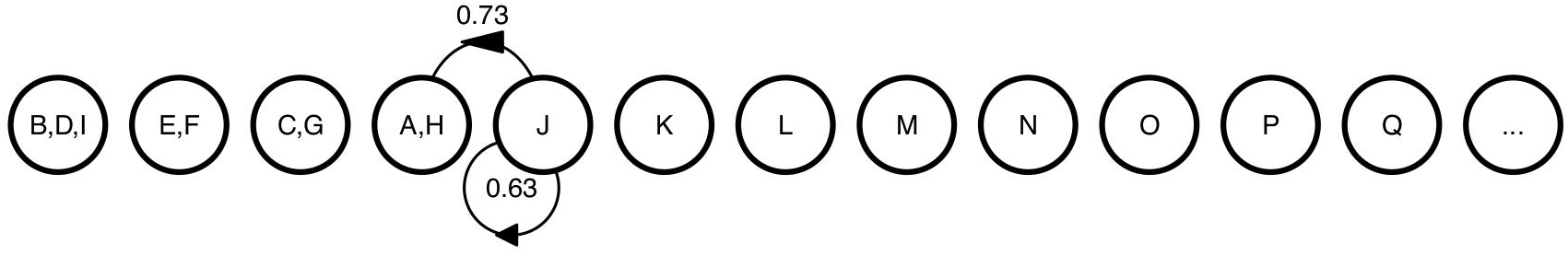

Again, by joining the OTU containing sequences A and H the number of TPs increases by 2 and the number of FNs decreases by 2 resulting in 8 TPs, 1210 TNs, 0 FPs, 7 FNs, and an improved MCC value of 0.73.

Processing sequence K is the same as for sequence A.

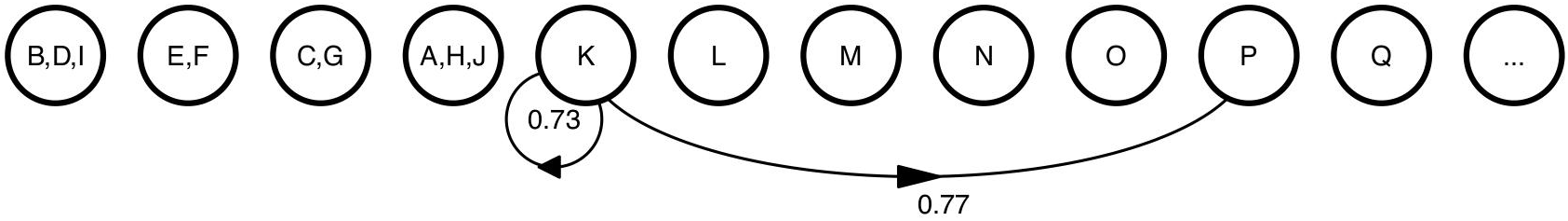

Merging sequences K and P into the same OTU results in 9 TPs, 1210 TNs, 0 FPs, 6 FNs, and an improved MCC value of 0.77.

Processing sequence L presents a more complicated set of decisions. Again, there are three choices. Because sequence L is similar to sequences O and P it could form an OTU with sequence O or with the OTU containing sequences K and P.

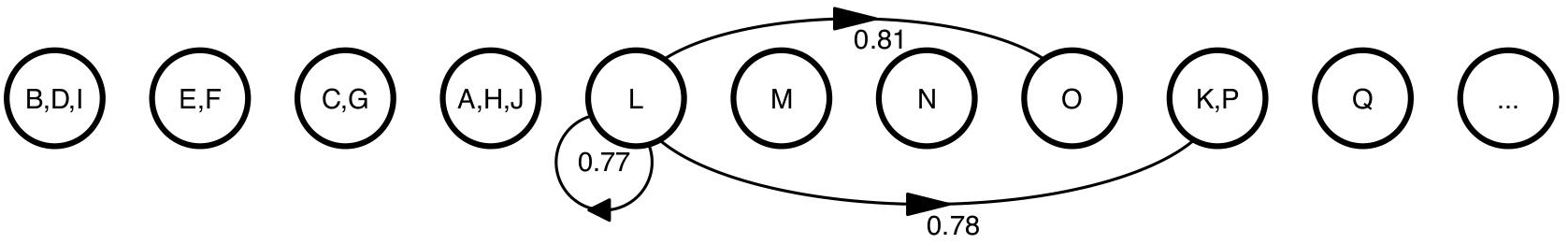

If sequence L remains on its own, the MCC value would remain 0.77. If it joined with sequence O the number of TPs would increase by 1 and the number of FNs would decrease by 1 resulting in an MCC value of 0.81. The subtlety of this step is found in when considering the possibility of sequence L joining an OTU with sequences K and P. It would increase the number of TPs by one and decrease the number of FN by one and by joining with sequence P; however, because O is not close to K the number of TNs would decrease by one and the number of FPs would increase by one. This would result in an MCC value of 0.78. Of the three options forming an OTU with sequences L and O provides the maximal MCC value. This results in 10 TPs, 1210 TNs, 0 FPs, 5 FNs, and an improved MCC value of 0.81.

The steps for processing sequence M is the same as earlier for sequence C.

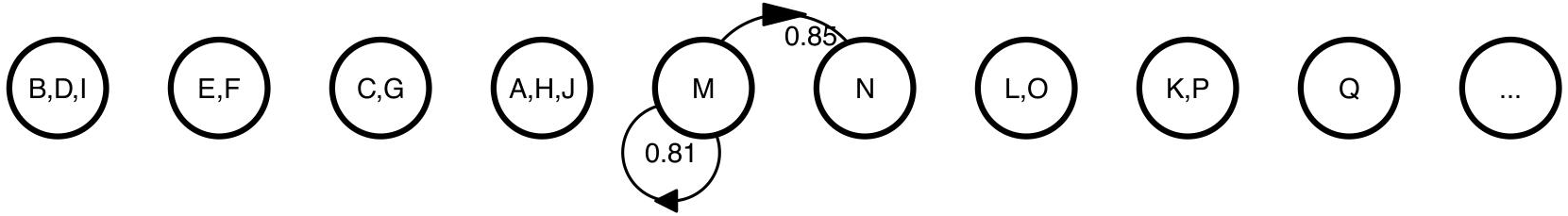

Merging sequences M and N into the same OTU results in 11 TPs, 1210 TNs, 0 FPs, 4 FNs, and an improved MCC value of 0.85.

Moving on to sequence N, the two options are to stay in an OTU with sequence M or to spit off and form a new OTU on its own.

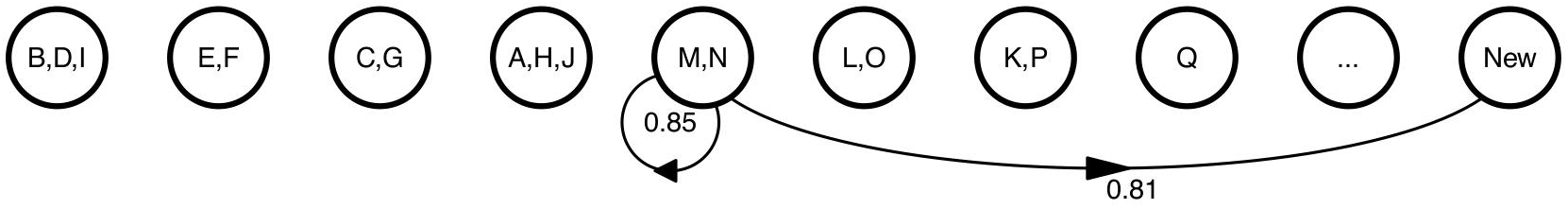

Remaining in an OTU with sequence M provides the larger MCC value and so the OTU memberships do not change.

Processing of sequence O presents three options that have been explored before. Sequence O can stay in its OTU with sequence L, it can join the OTU with sequences K and P, or it can form a new OTU on its own.

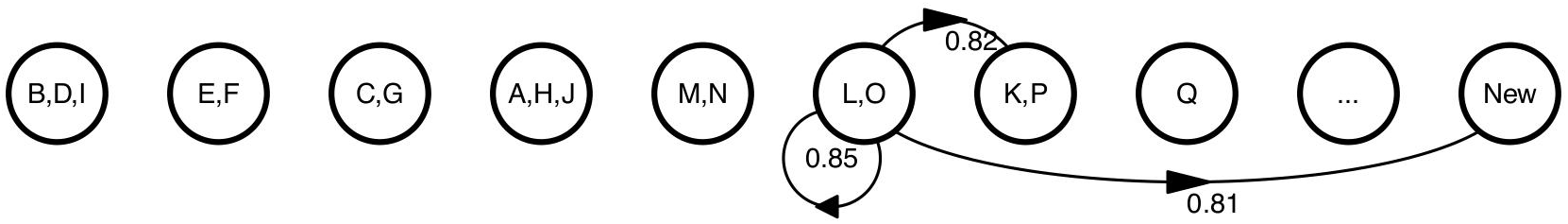

If sequence O remains in the OTU with sequence L, the MCC value would be 0.85. If it leaves that OTU to join sequences K and P in their OTU then the number of TNs would decrease by 1, but the number of FPs would increase by 1 because O is similar to P, but not to K. This would result in an MCC value of 0.82. If sequence O forms a new OTU on its own, then the number of TPs would decrease by one and the number of FNs would increase by one resulting in an MCC value of 0.81. The best option is for sequence O to remain in its OTU with sequence L.

For sequence P the steps taken are similar to those used to evaluate clusters for sequence O; however, the final decision is different. Sequence P can stay with sequence K in their OTU, it can leave to join the OTU with sequences L and O, or it can form a new OTU on its own.

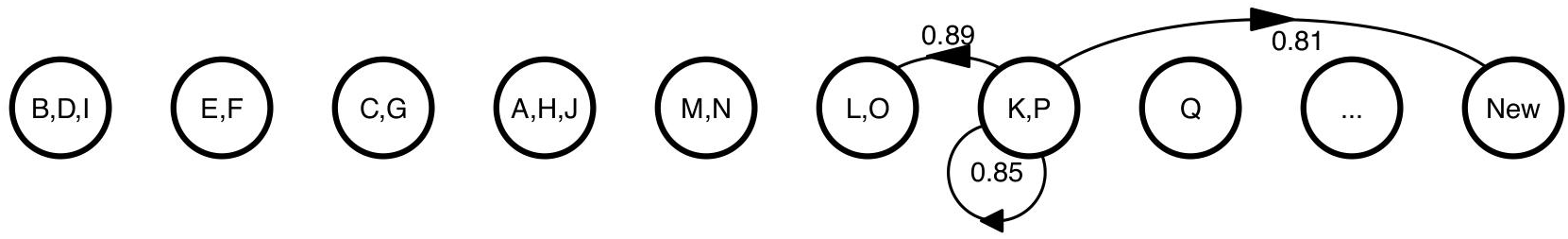

If sequence P remains in the OTU with sequence K, the MCC value would be 0.85. Alternatively, P could leave that OTU to join sequences L and O in their OTU. If P leaves its OTU with K then the number of TPs would decrease by one and the number of FNs would increase by one. By joining with L and O the number of TPs would increase by two and the number of FNs would decrease by two. The net effect would be to increase the number of TPs by 1 and decrease the number of FNs by 1. This would result in an MCC value of 0.89. If sequence P formed a new OTU on its own, then the number of TPs would decrease by one and the number of FNs would increase by one resulting in an MCC value of 0.81. The best option is for sequence P to leave its OTU with sequence K and join the OTU containing sequences L and O. The updated counts are 12 TPs, 1210 TNs, 0 FPs, 3 FNs.

To finish the first round of processing each sequence, sequence Q is processed. Sequence Q is similar to both sequences E and F. Because sequences E and F are in the same OTU, the situation is similar to processing sequence I.

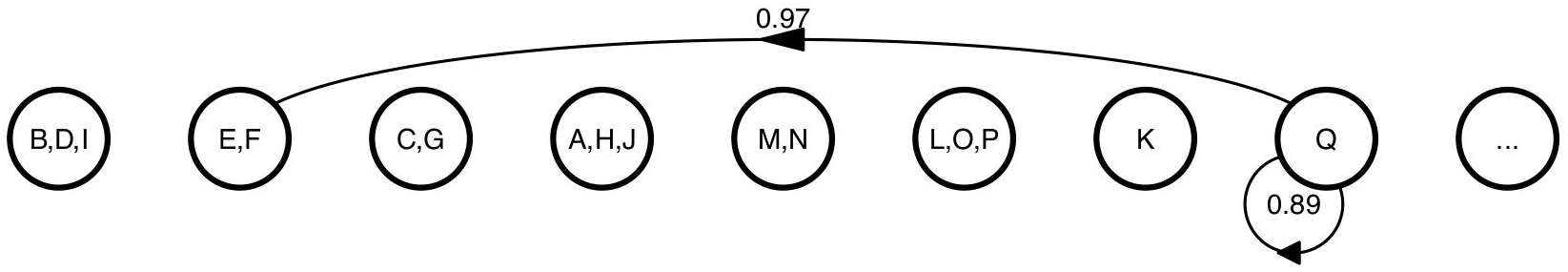

By joining the OTU containing sequences E and F the number of TPs increases by two and the number of FNs decreases by one. The updated counts are 14 TPs, 1210 TNs, 0 FPs, and 1 FNs, which result in an improved MCC value of 0.97.

Having processed each sequence, the first iteration of the algorithm is complete. The MCC value has changed from 0.00 to 0.97. Because the MCC value changed, it is necessary to re-evaluate each sequence again and re-evaluate the final MCC value to determine whether it has changed. In this case, evaluation of sequences A through J result in the same clustering pattern. When the algorithm reaches sequence K it finds that the sequence is similar to sequence P, which is in an OTU with L and O; however sequence K is not similar to L or O.

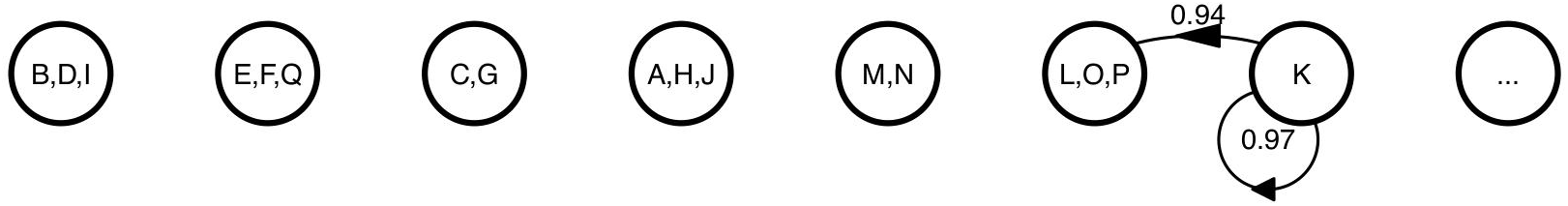

Although, sequence K is similar to sequence P, it is not similar to sequences L or O. Were sequence K to join their OTU, it would increase the number of TPs by one and decrease the number of FNs by one because of its similarity to sequence P, but it would increase the number of FPs by two and decrease the number of TNs by two because K is not similar to L or O. The end result would be a MCC value of 0.94, which is less than the MCC value of keeping sequence K on its own (i.e. 0.97).

Continuing the process for the remaining sequences, none of the sequences will move between OTUs and the MCC value does not change. At this point, the clustering has converged to the optimum MCC value of 0.97. Repeating this process using 100 different seeds for the random number generator required a median of 3 iterations (range from 2 to 4) to converge.

